## Supplementary figures and images for "Spatiotemporal Changes in Netrin/Dscam1 Signaling Dictate Axonal Projection Direction in *Drosophila* Small Ventral Lateral Clock Neurons"

### figure 1- figure supplement 1.pdf

**Figure 1-- figure supplement 1**

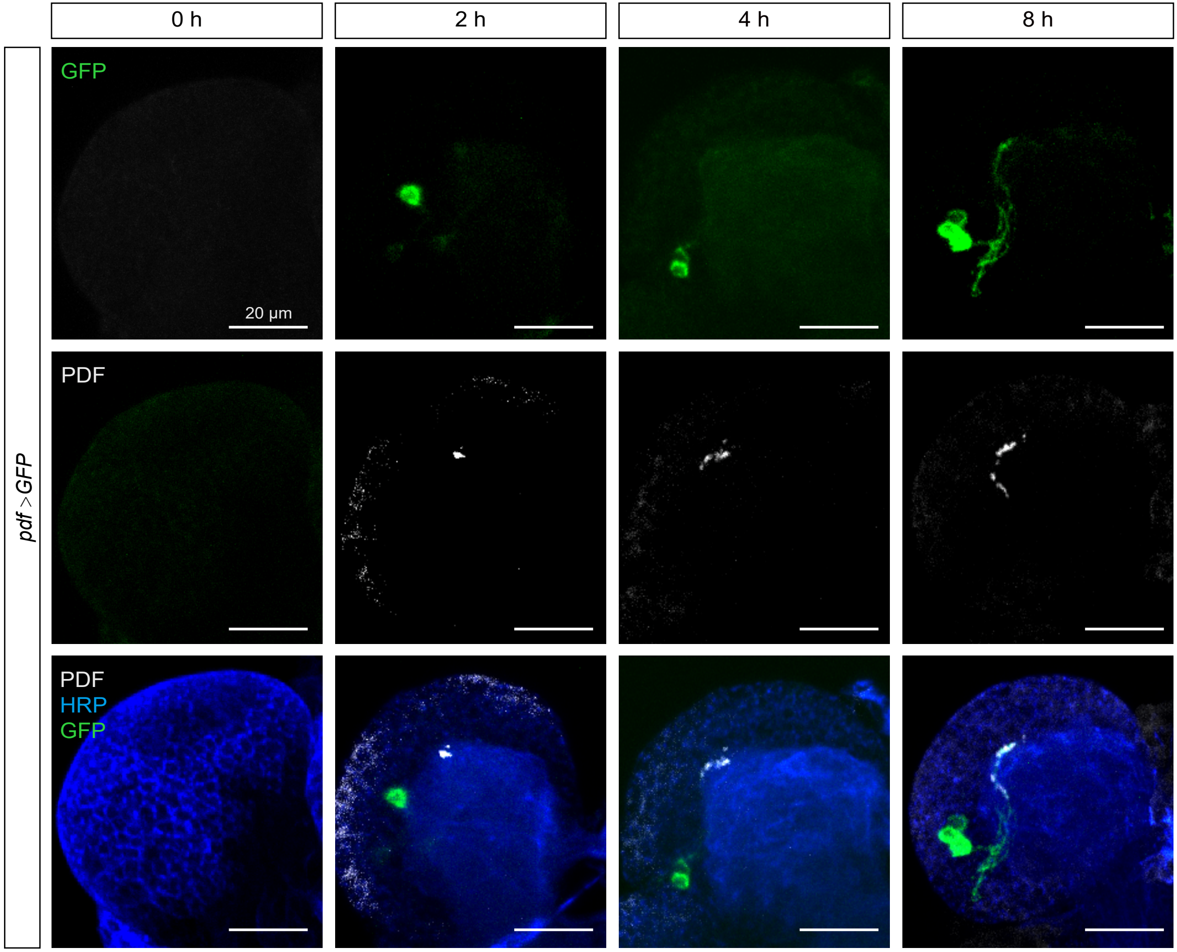

### figure 2- figure supplement 1.pdf

#### Figure 2-- figure supplement 1

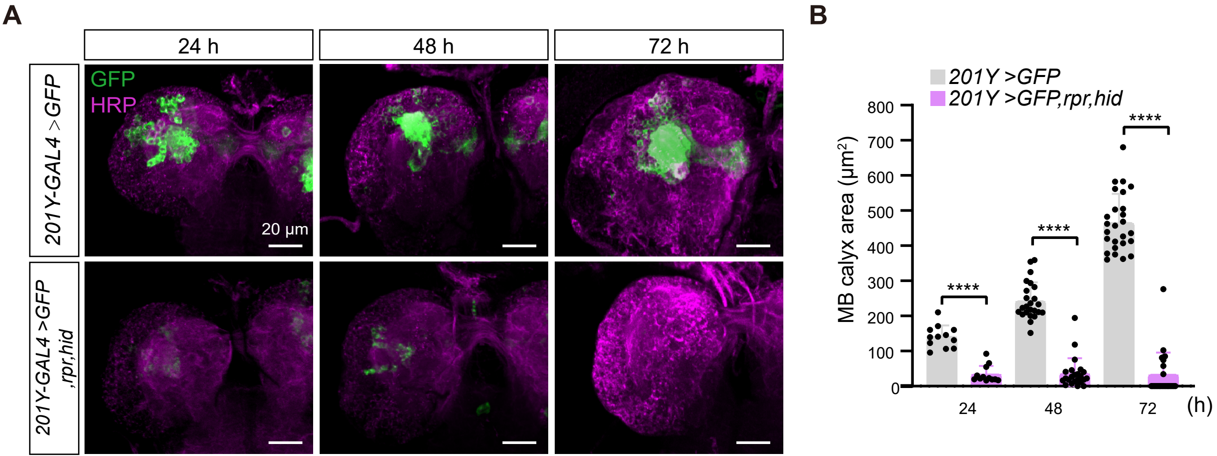

### figure 3- figure supplement 1.pdf

#### Figure 3-- figure supplement 1

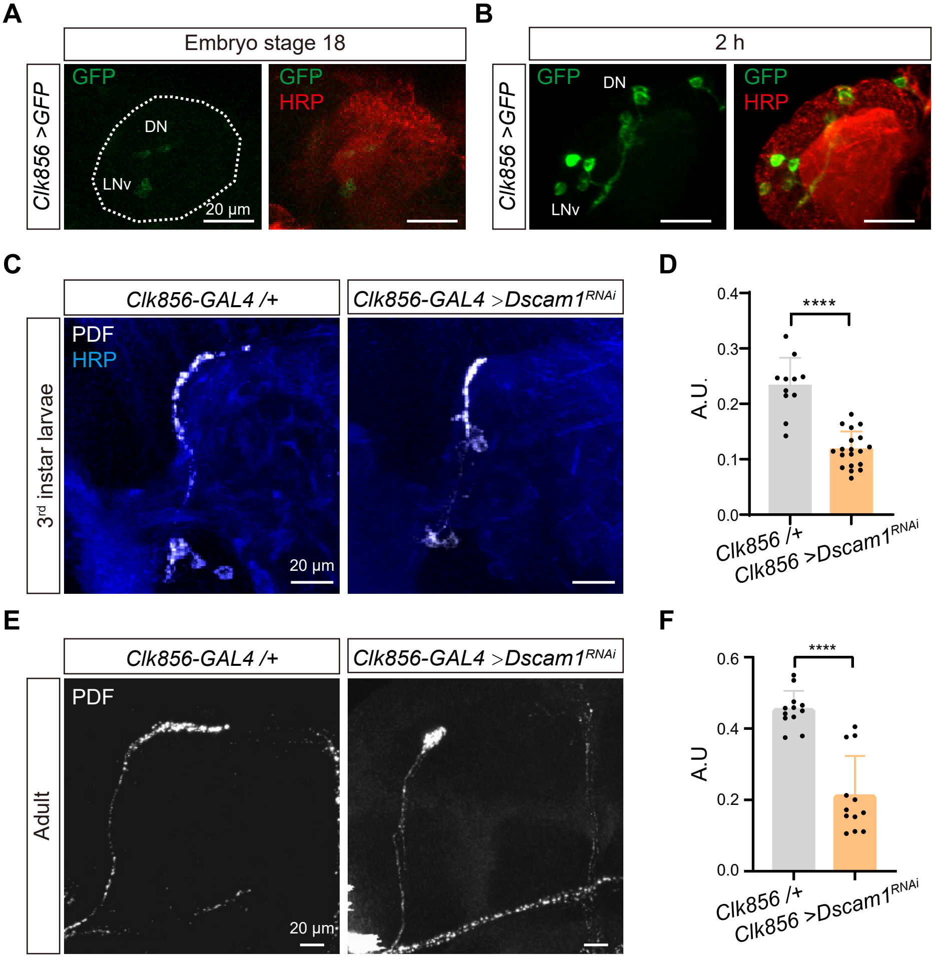

### figure 3- figure supplement 2.pdf

#### Figure 3-- figure supplement 2

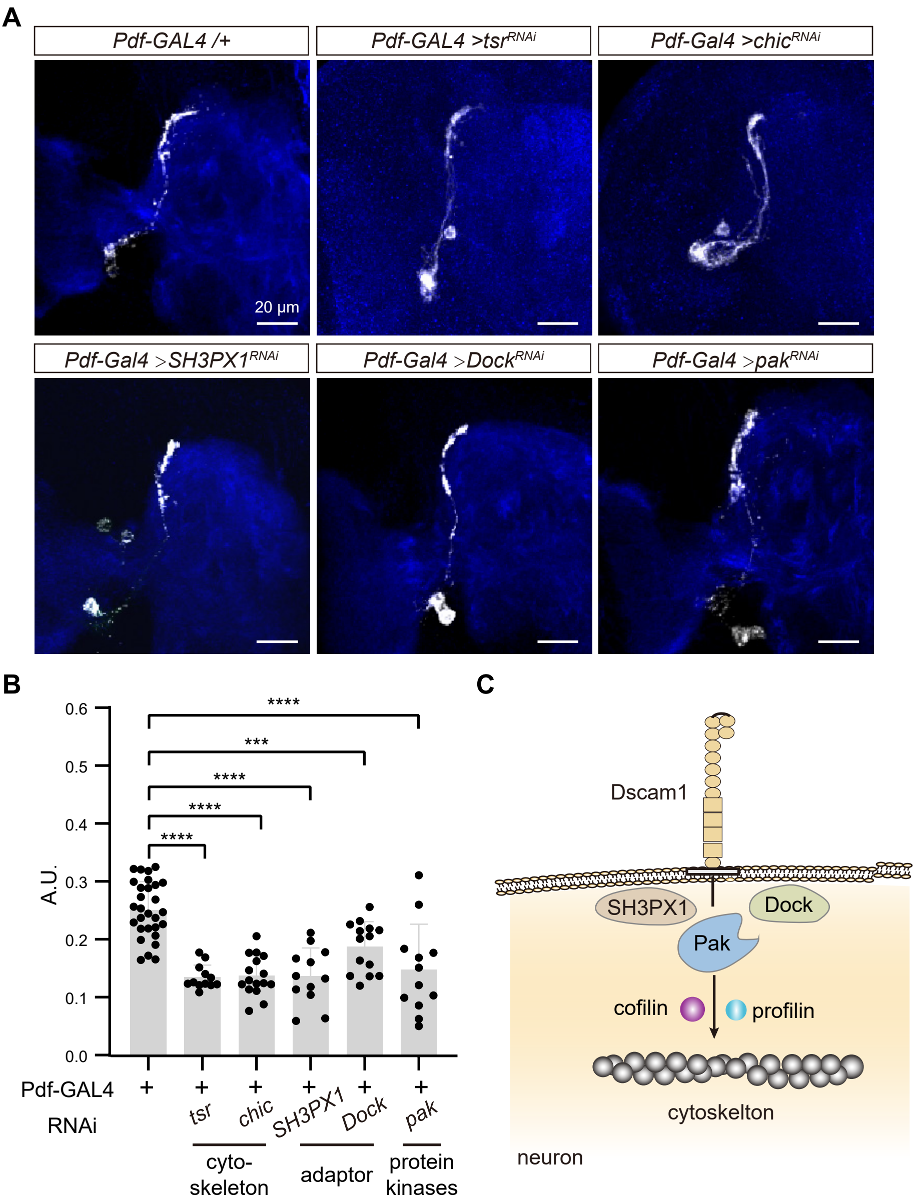

### figure 4- figure supplement 1.pdf

#### Figure 4-- figure supplement 1

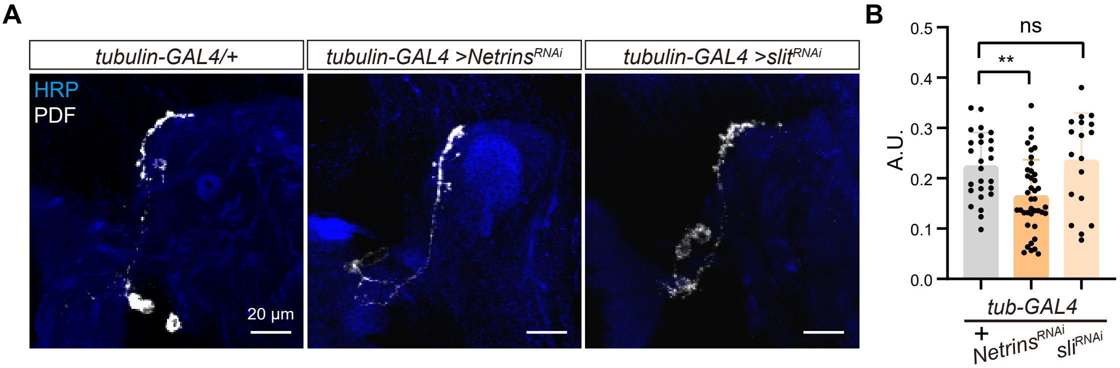

### figure 5- figure supplement 1.pdf

**Figure 5-- figure supplement 1**

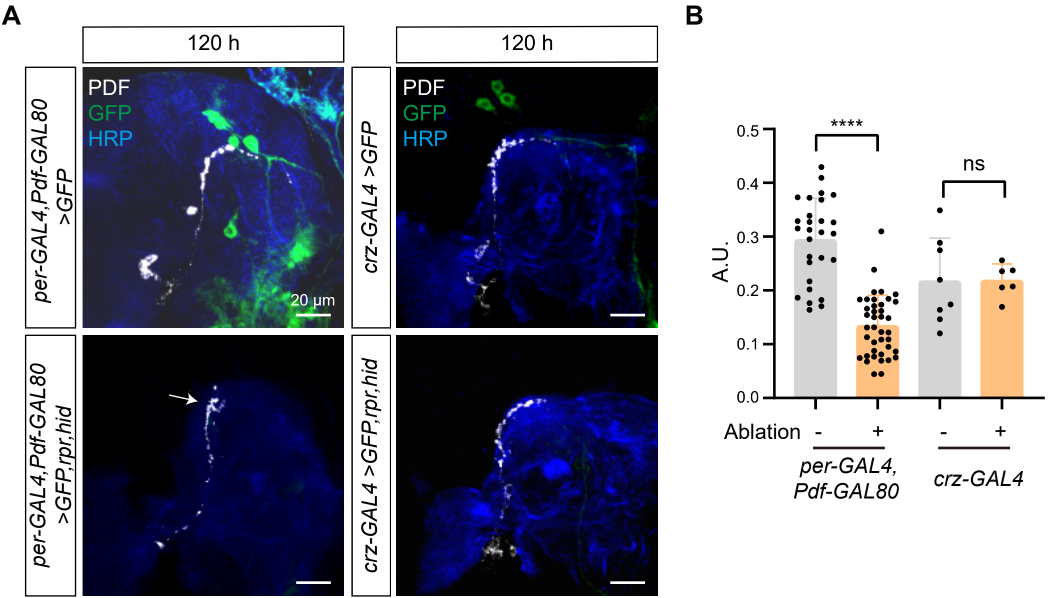

### figure 6- figure supplement 1.pdf

##### Figure 6-- figure supplement 1

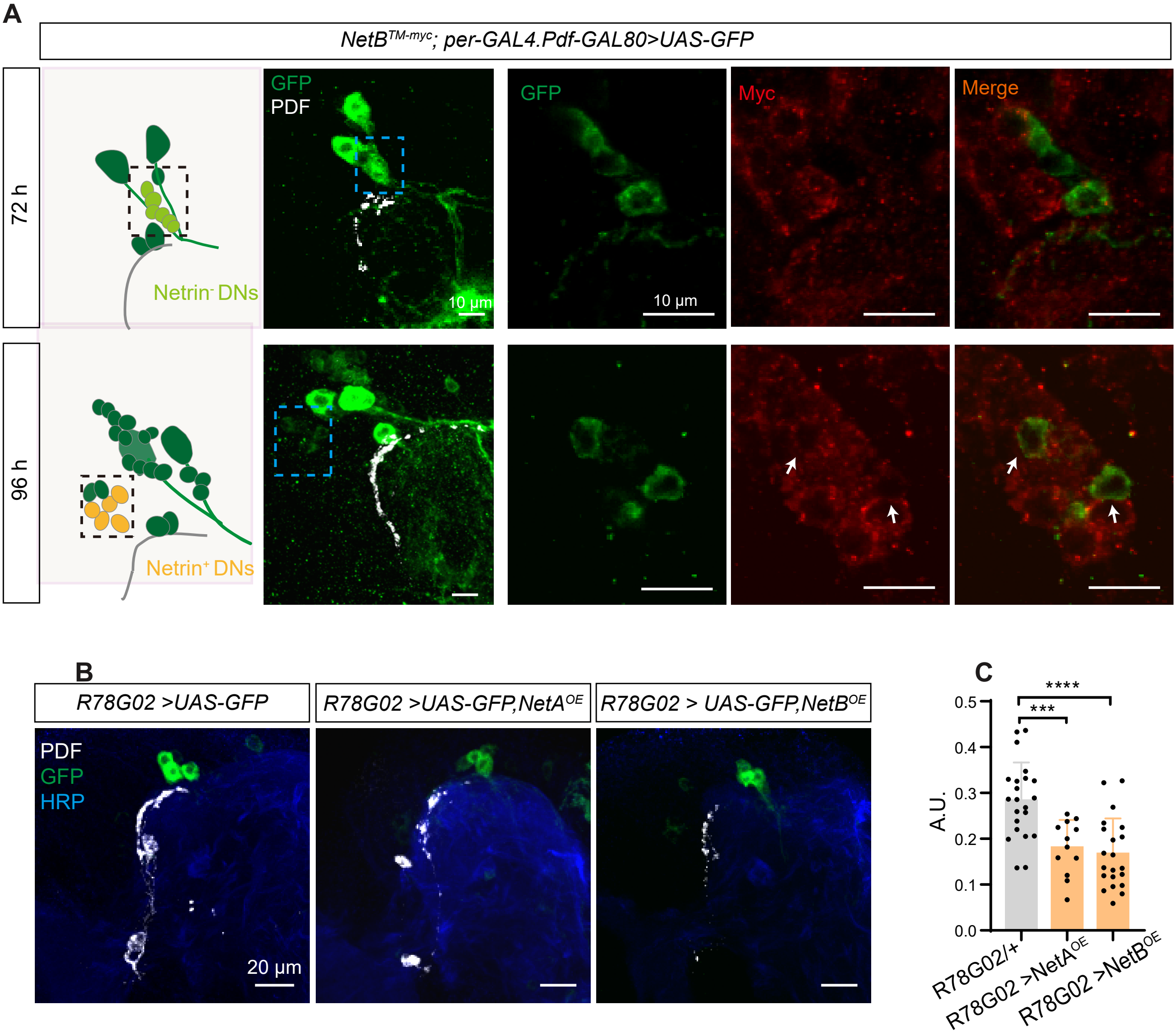
