## Supplementary File 1-3 for "Spatiotemporal Changes in Netrin/Dscam1 Signaling Dictate Axonal Projection Direction in *Drosophila* Small Ventral Lateral Clock Neurons": Supplementary File 1. Key resource table.docx

| **Key Resources Table** | | | | |
| --- | --- | --- | --- | --- |
| **Reagent type (species) or resource** | **Designation** | **Source or reference** | **Identifiers** | **Additional information** |
| genetic reagent (*D.*  *melanogaster*) | *Pdf-GAL4* | Bloomington Drosophila Stock Center | RRID: BDSC_6900 |  |
| genetic reagent (*D.*  *melanogaster*) | *nSyb-GAL4* | Bloomington Drosophila Stock Center | RRID: BDSC_51941 |  |
| genetic reagent (*D.*  *melanogaster*) | *repo-GAL4* | Bloomington Drosophila Stock Center | RRID: BDSC-7415 |  |
| genetic reagent (*D.*  *melanogaster*) | *OK107-GAL4* | Kyoto Stock Center | DGRC-106098 |  |
| genetic reagent (*D.*  *melanogaster*) | *Tab2-201Y-GAL4* | Bloomington Drosophila Stock Center | RRID:BDSC-4440 |  |
| genetic reagent (*D.*  *melanogaster*) | *Tubulin-GAL4* | Bloomington Drosophila Stock Center | RRID:BDSC-5138 |  |
| genetic reagent (*D.*  *melanogaster*) | *per-GAL4* | Bloomington Drosophila Stock Center | RRID:BDSC-7127 |  |
| genetic reagent (*D.*  *melanogaster*) | *MB247-GAL4* | Bloomington Drosophila Stock Center | RRID:BDSC-50742 |  |
| genetic reagent (*D.*  *melanogaster*) | *Crz-GAL4* | Bloomington Drosophila Stock Center | RRID:BDSC-51976 |  |
| genetic reagent (*D.*  *melanogaster*) | *Pdf-GAL80* | Bloomington Drosophila Stock Center | RRID:BDSC-80940 |  |
| genetic reagent (*D.*  *melanogaster*) | *UAS--mCD8-GFP* | Bloomington Drosophila Stock Center | RRID:BDSC-5137 |  |
| genetic reagent (*D.*  *melanogaster*) | *UAS--mCD8-GFP* | Bloomington Drosophila Stock Center | RRID:BDSC-5130 |  |
| genetic reagent (*D.*  *melanogaster*) | *UAS--mCD8-RFP* | Bloomington Drosophila Stock Center | RRID:BDSC-27392 |  |
| genetic reagent (*D.*  *melanogaster*) | *lexAop-mCD8-GFP* | Bloomington Drosophila Stock Center | RRID: BDSC-32207 |  |
| genetic reagent (*D.*  *melanogaster*) | *Clk856-GAL4* | Bloomington Drosophila Stock Center | RRID:BDSC-93198 |  |
| genetic reagent (*D.*  *melanogaster*) | *UAS-rprC;;UAS-hid* | Gifted from Yufeng Pan | N/A |  |
| genetic reagent (*D.*  *melanogaster*) | *Dscam1 ^1^* | Bloomington Drosophila Stock Center | RRID:BDSC-5934 |  |
| genetic reagent (*D.*  *melanogaster*) | *Dscam1 ^21^* | Gifted from Haihuai He | N/A |  |
| genetic reagent (*D.*  *melanogaster*) | *Dscam1 ^05518^* | Bloomington Drosophila Stock Center | RRID:BDSC-11412 |  |
| genetic reagent (*D.*  *melanogaster*) | *NetA^Δ^* | Bloomington Drosophila Stock Center | RRID:BDSC-66878 |  |
| genetic reagent (*D.*  *melanogaster*) | *NetB^Δ^* | Bloomington Drosophila Stock Center | RRID:BDSC-66879 |  |
| genetic reagent (*D.*  *melanogaster*) | *NetAB^Δ^* | Bloomington Drosophila Stock Center | RRID:BDSC-66877 |  |
| genetic reagent (*D.*  *melanogaster*) | *NetB-GFP* | Bloomington Drosophila Stock Center | RRID:BDSC-67644 |  |
| genetic reagent (*D.*  *melanogaster*) | *NetB^tm^* | Bloomington Drosophila Stock Center | RRID:BDSC-66880 |  |
| genetic reagent (*D.*  *melanogaster*) | *UAS-Dscam1^RNAi^* | Tsinghua Fly Center | THU3896 |  |
| genetic reagent (*D.*  *melanogaster*) | *UAS-NetA^RNAi^* | Tsinghua Fly Center | THU1972 |  |
| genetic reagent (*D.*  *melanogaster*) | *UAS-NetB^RNAi^* | Tsinghua Fly Center | TH201500623.S |  |
| genetic reagent (*D.*  *melanogaster*) | *UAS-slit^RNAi^* | Tsinghua Fly Center | THU1910 |  |
| genetic reagent (*D.*  *melanogaster*) | *UAS-pak^RNAi^* | Tsinghua Fly Center | TH201500668.S |  |
| genetic reagent (*D.*  *melanogaster*) | *UAS-Dock^RNAi^* | Tsinghua Fly Center | THU2815 |  |
| genetic reagent (*D.*  *melanogaster*) | *UAS-SH3PX1^RNAi^* | Tsinghua Fly Center | THU2738 |  |
| genetic reagent (*D.*  *melanogaster*) | *UAS-tsr^RNAi^* | Tsinghua Fly Center | THU0972 |  |
| genetic reagent (*D.*  *melanogaster*) | *UAS-chic^RNAi^* | Tsinghua Fly Center | THU0986 |  |
| antibody | anti-PDF mouse monoclonal antibody | DSHB | C7; RRID: AB_760350 | IF(1:300) |
| antibody | anti-HRP rabbit monoclonal antibody | Jackson Immuno Research | Cat# 323-005-021, RRID:AB_2314648 | IF(1:500) |
| antibody | anti-GFP rabbit polyclonal FLUR^TM^488 | Invirtrogen | Cat# A-21311, RRID:AB_221477 | IF(1:200) |
| antibody | anti-RFP polyclonal rabbit antibody | Rockland | Cat# 600-401-379, RRID:AB_2209751 | IF(1:500) |
| antibody | anti-GFP polyclonal chicken antibody | Invirtrogen | Cat# A10262, RRID:AB_2534023 | IF(1:2000) |
| antibody | anti-Myc polyclonal Rabbit antibody | Cell Signaling Technology | Cat# 9402, RRID:AB_2151827 | IF(1:200) |
| antibody | Anti-Dscam1 18 mAb mouse monoclonal antibody | Gift from Tzumin Lee (Yu et al., 2009) | N/A | IF(1:20) |
| antibody | Alexa Fluor 555 goat anti-rabbit IgG, | Abcam | Cat# ab150078; RRID: AB_2722519 | IF(1:200) |
| antibody | Alexa Fluor 647 goat anti-mouse IgG | Abcam | Cat# ab150115, RRID:AB_2687948 | IF(1:200) |
| antibody | Alexa Fluor 488 goat anti-rabbit IgG | Abcam | Cat# ab150081; RRID: AB_2734747 | IF(1:200) |
| antibody | Alexa Fluor 488 goat anti-mouse IgG | Abcam | Cat#ab150113; RRID:AB_2576208 | IF(1:200) |
| antibody | Alexa Fluor 488 goat anti-chicken IgY | Abcam | Cat# A-11039, RRID:AB_2534096 | IF(1:200) |
| recombinant DNA reagent | pUAST-HA-NetA (plasmid) | Gift from Duan R | N/A |  |
| recombinant DNA reagent | pUAST-HA-NetB (plasmid) | Gift from Duan R | N/A |  |
| software, algorithm | GraphPad Prism 8.0.2 | GraphPad | RRID: SCR_002798 |  |
| software, algorithm | Zeiss LSM Image Browser | Zeiss | https://www.zeiss.com/microscopy/int/downloads/lsm-5-series.html |  |
| software, algorithm | fiji | Image J | RRID: SCR_002285 |  |
